# Influenza A Virus infection is associated with TDP-43 pathology and neuronal damage in the brain

**DOI:** 10.1101/2025.06.30.662477

**Authors:** Min-Tae Jeon, Dong-Hwi Kim, Sinnead A Cogill, Kyusung Kim, In Yeong Park, Jae-Hyeong Kim, Minyeop Nam, Do-Geun Kim, In-Soo Choi

## Abstract

Viral pandemics such as COVID-19 have demonstrated long-term neurological consequences, including memory impairment and depression, emphasizing the importance of understanding virus–brain interactions [1]. Similar concerns have been raised for Influenza A virus (IAV), which has been implicated in neurodegenerative disorders [2, 3]. In this study, we investigated the neuropathological effects of highly pathogenic avian influenza (HPAI) H5N1 and H5N8 strains in a mouse model. Although viral RNA was detected in the brain post-infection, no viral proteins were found, suggesting limited or transient brain replication. Despite this, infected brains showed significant neuronal damage, including axonal loss and nuclear condensation, as evidenced by immunofluorescence and Nissl staining. We also observed pathological changes in TDP-43, including conformational alterations and increased phosphorylation, which required antigen retrieval for detection—features reminiscent of those found in frontotemporal dementia and amyotrophic lateral sclerosis [4, 5]. Transcriptomic analysis further revealed strain-specific host responses, including activation of interferon-related genes and downregulation of microtubule-associated pathways. These findings suggest that IAV infection can trigger hallmarks of neurodegeneration in the absence of persistent viral protein expression, possibly through host-driven mechanisms. Our results underscore the need for further investigation into virus-induced molecular pathways contributing to neurodegenerative disease.

## Results and Discussion

### Neurodegenerative features in influenza A virus–infected mouse brains

To examine whether highly pathogenic avian influenza (HPAI) infection elicits neuropathological changes in the brain, we intranasally infected BALB/c mice with either the H5N1 or H5N8 strain of Influenza A virus (IAV), both of which are of avian origin and associated with pandemic potential due to their capacity for cross-species transmission and genomic reassortment [2, 3]. Although IAV is primarily a respiratory pathogen, several studies have reported its ability to invade the central nervous system (CNS), with viral RNA detected in cerebrospinal fluid and brain tissue in both experimental models and clinical cases [6-9]. Mice were sacrificed at six days post-infection (dpi), a time point selected based on our previous report showing robust pulmonary inflammation, cytokine induction, and histopathological damage in the lung at this stage in the same model [10], confirming the establishment of systemic viral infection.

To assess CNS involvement, we performed RT-PCR analysis targeting the IAV M gene and detected viral RNA in brain tissue from both H5N1- and H5N8-infected animals (Fig. 1A), indicating viral dissemination beyond the respiratory tract. However, despite the presence of viral RNA, immunostaining with multiple antibodies failed to detect viral proteins in the brain (Fig. S1A, B). In contrast, LPAI H1N1-infected animals displayed clear neuronal expression of viral proteins (Fig. S1C), suggesting that while both viral types reach the brain, HPAI may differ in replication efficiency or antigen stability within the CNS. The inability to detect viral proteins could also reflect rapid clearance, limited translation, or host restriction mechanisms active in neural tissue.

**Figure 1.**
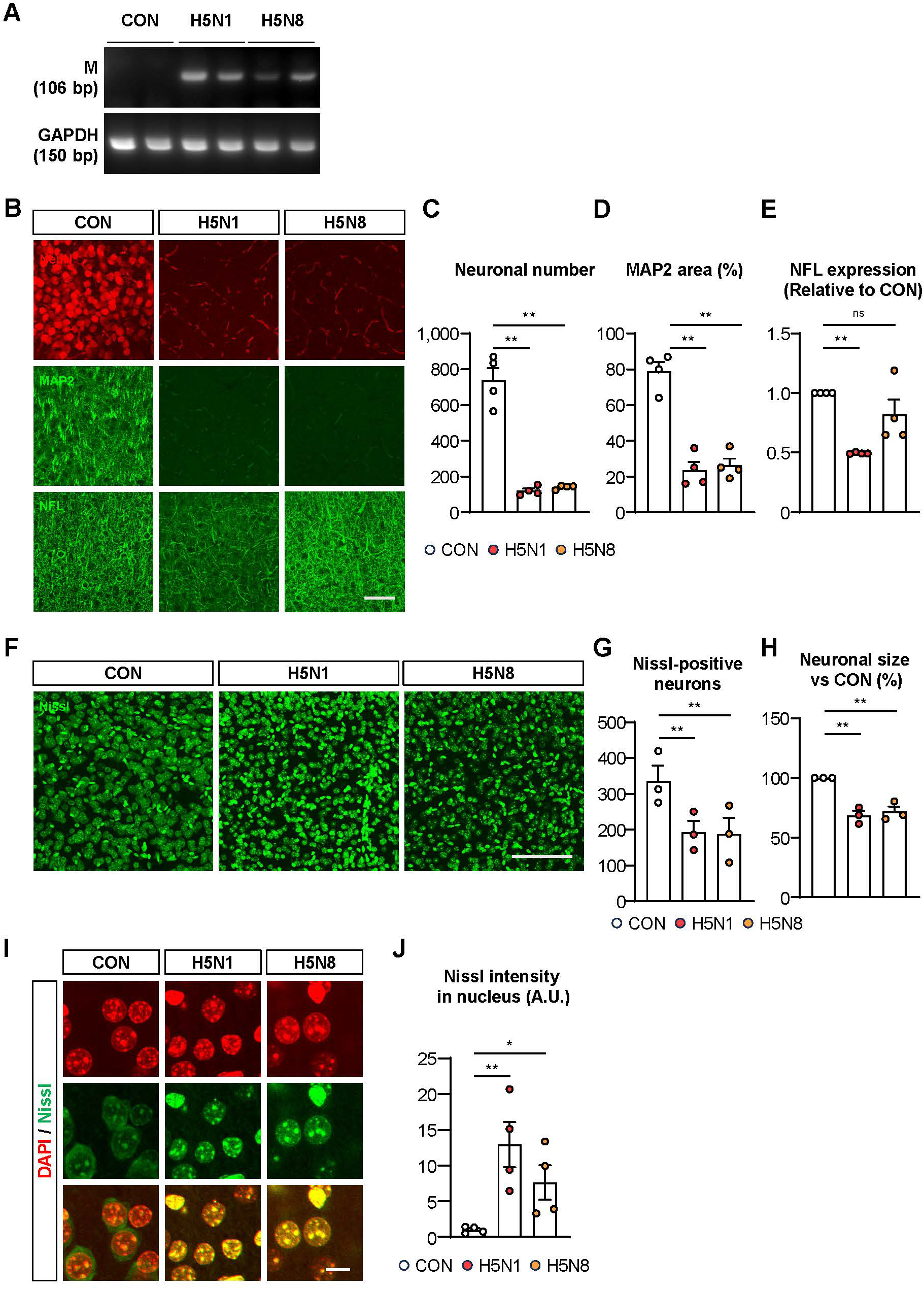
Influenza A viral infection in mouse brain leads to an increase in neuronal damage. **A**. PCR analysis revealed the presence of viral RNA in the mouse brain, indicating that influenza A virus can invade the brain. **B**. Immunostaining showing neuronal activity through NeuN (upper; red), MAP2 (middle; green) and NFL (lower; green). Scale bar, 100 μm. **C-E**. Graphs indicate NeuN-positive neuronal number (C), MAP2 expression area (D) and NFL expression (E) in the HPAI-infected mouse cortex. (^**^ *p*< 0.01, determined by one-way analysis of variance (ANOVA) and Dunnet’s multiple comparison’s test, error bars indicate means with SEM; n = 4). **F**. Nissl staining with NeuroTracer (green) shows RNA coverage in neurons. Scale bar, 100 μm. **G and H**. Graphs indicate Nissl-positive neuron count (G) and Nissl-positive neuronal size (H). **I**. Nissl staining with NeuroTracer (green) shows RNA coverage in brain tissue. Scale bar,10 μm. J. Graph indicates Nissl-positive neuron intensity in nucleus (n=4). All histograms shows quantitative values as mean with SEM (^*^p < 0.05 and ^**^p < 0.01, determined by one-way analysis of variance (ANOVA) and Dunnet’s multiple comparison’s test).

To determine whether viral entry is associated with blood-brain barrier (BBB) dysfunction, we assessed key markers of BBB integrity. Immunostaining for endothelial tight junction proteins and astrocyte–endfoot markers revealed no detectable structural abnormalities in the infected groups compared to controls (Fig. S2A–D), suggesting that IAV RNA enters the CNS without overt barrier disruption. This is in contrast to other neurotropic viruses, which commonly disrupt BBB architecture [11, 12].

We next evaluated glial responses to infection. Astrocyte activation, as assessed by ALDH1L1 and S100-β immunostaining, was not observed in HPAI-infected animals (Fig. S3A), indicating a lack of widespread astrogliosis. In contrast, Iba1-positive microglia exhibited activated morphologies and increased density in the cortex of H5N1-infected animals (Fig. S3B), consistent with previous reports linking IAV to microglial-mediated neuroinflammation and cytokine production [13]. These results suggest that microglial activation may precede or occur independently of astrocytic responses and that strain-specific differences in neuroimmune activation exist.

To investigate neuronal integrity, we performed immunofluorescence staining for NeuN, MAP2, and NFL, which respectively label neuronal soma, dendrites, and axons. All three markers exhibited reduced signal intensity in H5N1- and H5N8-infected brains relative to controls (Fig. 1B–1E), indicating neuronal structural loss across multiple compartments. To further assess neuronal viability, Nissl staining was performed and revealed a decrease in both the number and size of Nissl-positive neurons (Fig. 1F–1H), consistent with neuronal loss and shrinkage. Notably, Nissl staining also showed increased signal intensity within neuronal nuclei in infected groups (Fig. 1I, J). Given that Nissl dyes bind ribosomal RNA and are typically confined to the cytoplasm, this abnormal nuclear staining pattern may reflect disrupted RNA trafficking or nucleolar stress, as has been reported in early stages of neurodegenerative disease models [14].

### TDP-43 pathology and strain-specific molecular responses following IAV infection

Given the observed nuclear retention of Nissl signal and loss of cytoplasmic RNA integrity, we next investigated the behavior of TDP-43 (TAR DNA-binding protein 43), a nuclear RNA-binding protein with established roles in RNA splicing, mRNA trafficking, and microRNA biogenesis [15, 16]. TDP-43 is known to undergo pathological mislocalization and phosphorylation in several neurodegenerative diseases, including amyotrophic lateral sclerosis (ALS) and frontotemporal dementia (FTD), where it forms insoluble cytoplasmic inclusions and is depleted from the nucleus [4, 5].

Western blot analysis of cortical lysates from HPAI-infected mice revealed a marked increase in total TDP-43 protein levels compared to controls (Fig. 2A). However, immunofluorescence failed to detect TDP-43 in the infected brains under native staining conditions, while strong nuclear TDP-43 signal was observed in uninfected animals (Fig. 2B). TDP-43 signal in the infected brain became detectable only after antigen retrieval, suggesting that conformational changes—possibly associated with protein aggregation or epitope masking—had occurred, as similarly described in human pathological specimens from ALS patients [4]. Furthermore, immunoblotting and immunostaining showed elevated levels of phosphorylated TDP-43 (pTDP-43) in infected brains (Fig. 2C, D), a pathological post-translational modification that contributes to the formation of insoluble aggregates and nuclear clearance [5]. These findings indicate that IAV infection induces biochemical and structural alterations in TDP-43 that resemble those observed in TDP-43 proteinopathies. Importantly, immunostaining for Alzheimer’s disease–related proteins, including amyloid-β and phospho-Tau, showed no detectable signal in infected brains (Fig. S5), suggesting that the degenerative phenotype is not associated with classical AD pathology [17].

**Figure 2.**
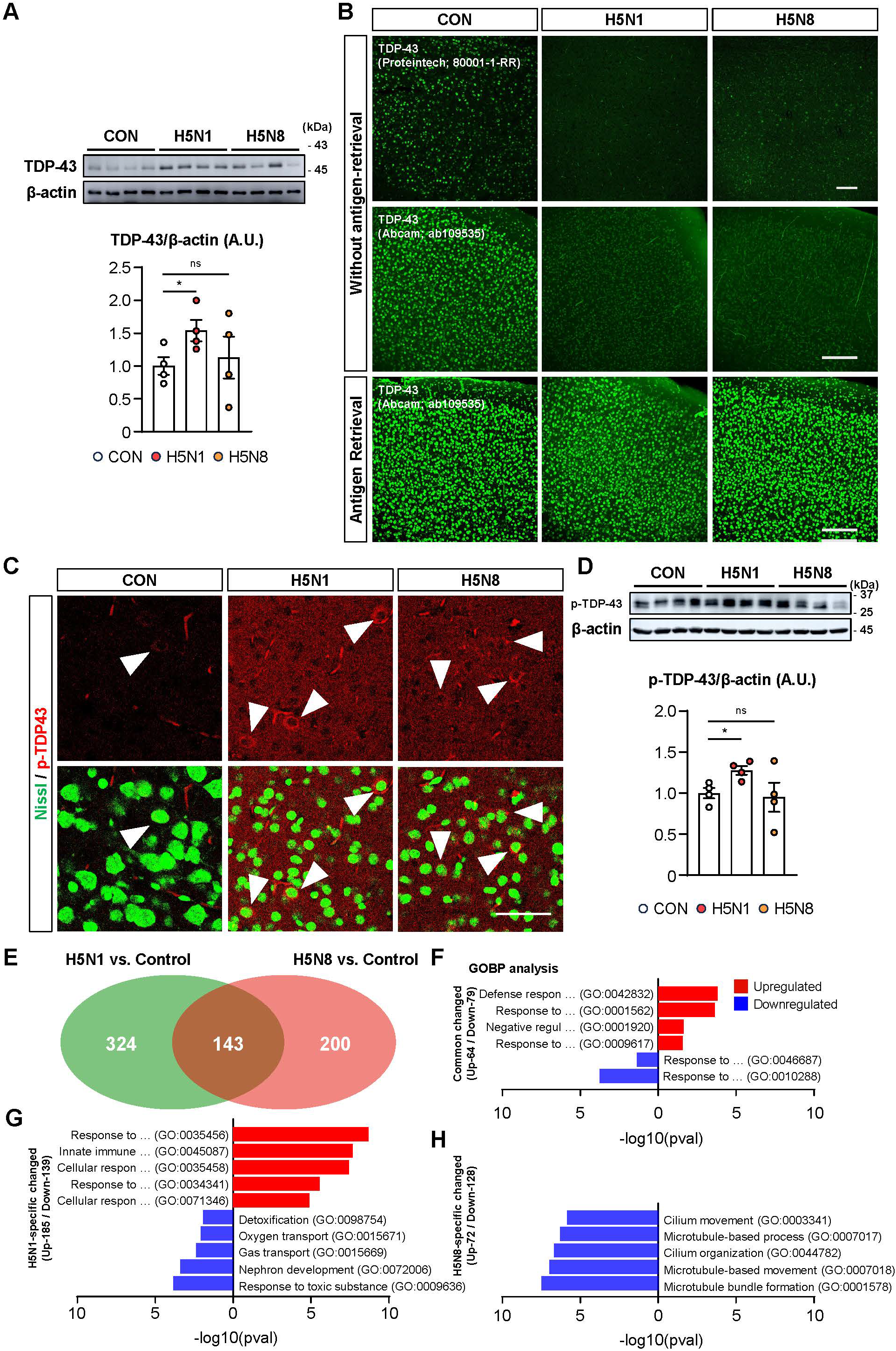
Pathological changes in the brain following Influenza A infection. **A**. Western blotting results for TDP-43 protein in brain tissue for three treatment groups (n = 4). **B**. Brain sections were immunostained with TDP-43 antibody to visualize protein depletion in infected brain tissue following antigen retrieval. Scale bars, 100 μm. **C**. Immunostaining for phosphorylated TDP-43 (red). Scale bars, 50 μm. **D**. Western blotting results for p-TDP-43 protein in brain tissue for three treatment groups (n = 4). **E-H**. RNA sequencing reveals strain-specific mechanisms of immune regulation. **E**. The Venn diagram represents the number of changed genes between the control and viral-infected groups. **F-H**. Histograms illustrate the highly enriched Gene Ontology Biological Process (GOBP) terms associated with significantly altered genes identified in D.

To investigate whether the observed neurodegenerative features are accompanied by broader transcriptional changes, we performed RNA sequencing (RNA-seq) of cortical tissue from H5N1- and H5N8-infected mice. Principal component analysis revealed clear segregation between infected and control samples, indicating a robust virus-induced transcriptional response (Fig. S6A, B). Differential expression analysis showed that H5N1-infected brains had 185 upregulated and 139 downregulated genes, while H5N8-infected brains exhibited 72 upregulated and 128 downregulated genes (Fig. S6C). A direct comparison between the two infected groups highlighted both shared and strain-specific expression patterns (Fig. 2E).

Gene ontology (GO) enrichment analysis of shared upregulated genes in both infection groups revealed significant enrichment in terms related to antiviral defense, innate immune activation, and cellular stress responses (Fig. 2F). Notably, genes specifically upregulated in H5N1-infected brains were strongly associated with type I interferon signaling, cytokine activity, and host defense (Fig. 2G), consistent with the more pronounced microglial activation observed histologically. In contrast, H5N8-infected brains exhibited fewer significantly enriched GO terms among upregulated genes but showed enrichment for downregulated pathways related to microtubule organization, cytoskeletal stability, and axonal transport (Fig. 2H). These differences suggest that while both strains trigger innate immune responses, H5N1 induces a more inflammatory transcriptional signature, whereas H5N8 preferentially downregulates structural pathways involved in neuronal maintenance [18].

Taken together, these results suggest that HPAI infection disrupts neuronal homeostasis through strain-specific molecular pathways and induces features reminiscent of TDP-43 proteinopathy. Importantly, these effects occurred in the absence of detectable viral protein in the brain, implying that persistent infection is not required to drive neuropathological outcomes. While we cannot conclusively establish a causal link between TDP-43 alterations and neuronal death in this study, the temporal and spatial co-occurrence of pTDP-43 accumulation, RNA retention, and axonal damage raises the possibility of a common mechanism of neurotoxicity. Similar TDP-43 abnormalities have been observed in viral infections including SARS-CoV-2 and coxsackievirus B3, where viral proteins modulate host RNA-binding proteins to impair neuronal function [19-21]. These parallels suggest that IAV, despite its predominantly respiratory tropism, may also contribute to neurodegenerative changes via host RNA-processing dysfunctions. Further mechanistic studies will be required to define the upstream triggers of TDP-43 pathology in the context of acute viral infection.

## Materials and Methods

### Virus strains, animals and ethics statement

One subtype of low pathogenic avian influenza (H1N1; A/PR/8/34) and two subtypes of HPAI (H5N1 clade 2.3.2.1; A/chicken/Korea/Gimje/2008 and H5N8 clade 2.3.4.4b; A/duck/Korea/H338/2020) were used to infect 7-8 week-old BALB/c mice (Orient Bio, South Korea). Mice were housed in a biosafety level 3 facility at Konkuk University with controlled temperature, humidity, and lighting. After 1 week of acclimation, mice were intranasally infected with 1 × 10^2^ PFU of each virus or PBS. Six days post-infection, the mice were sacrificed, and their brains were collected, washed with PBS, and fixed in 10% neutral buffered formalin for 48 hours. This study was approved by Konkuk University’s Animal Research Center (Approval No. KU23140) and Institutional Biosafety Committee (Accreditation No. KUIBC-2023-16).

### Immunohistochemistry

Half brains were washed with PBS and fixed in 4% paraformaldehyde (PFA) for 48 hours. Brain sections (50 μm) were used for immunofluorescence staining. Sections were permeabilized with 1% Triton-X in PBS for 20 min, blocked with 5% NGS/PBS for 45 min, and incubated overnight with primary antibodies at 4^°^C. After rinsing, sections were incubated with fluorescent secondary antibodies for 1 hour at room temperature. Sections were mounted with DAPI Fluoromount-G. For TDP-43 staining, antigen retrieval was performed by heating sections in PBS (100^°^C) for 20-40 minutes before antibody staining. Images were acquired with a confocal microscope (Leica) and analyzed with LAS X software.

To assess neuronal damage in the cortex, brain sections were stained with various markers respectively. Images was quantified in 3 rectangular boxes (0.5×0.3 mm) on the cortical layer V for each section.

### Western Blotting

Half brains were incubated for viral inactivation at 56^°^C for 30 minutes and treated with RNAlater (ThermoFisher) for protein and RNA analysis. Brain homogenates were lysed in RIPA buffer with protease and phosphatase inhibitors. Protein concentrations were determined using the Pierce BCA Protein Assay kit (ThermoFisher). Equal protein amounts were separated on SDS-PAGE gels, transferred to PVDF membranes, and blocked with 5% BSA/TBS-T. Membranes were incubated overnight with primary antibodies. After washing, membranes were incubated with HRP-conjugated secondary antibodies (Jackson ImmunoResearch) for 1 hour, and protein complexes were detected using Lumigen chemiluminescence reagent (Lumigen) on the LAS-500 imager (GE Healthcare).

### RNA sequencing

Total RNA concentration was measured using the Quant-IT RiboGreen assay (Invitrogen). RNA integrity was assessed with the TapeStation RNA screentape (Agilent), selecting samples with a RIN > 7.0 for library preparation. For library construction, 0.5 μg of RNA was processed using the Illumina TruSeq Stranded Total RNA Library Prep Gold Kit. rRNA was removed using the Ribo-Zero rRNA Removal Kit (Illumina), and RNA was fragmented with divalent cations at elevated temperatures. First-strand cDNA was synthesized with SuperScript II reverse transcriptase, followed by second-strand cDNA synthesis and end repair. The final library was amplified, quantified using KAPA Library Quantification kits, and quality-checked with TapeStation D1000. Indexed libraries were sequenced (2×151 bp) on the Illumina NovaSeq 6000 by Macrogen.

### Statistical analysis

More than 3 animals for each group were measured in this study. The precious number of animals are given in the figure legends. Differences between groups were analyzed using one-way ANOVA with Dunnett’s multiple comparison test. All analyses were performed using GraphPad Prism 8 (GraphPad Software, San Diego, CA).

## Acknowledgements

This research was supported by the Konkuk University Researcher Fund in 2024 (grant number: 2024-A019-0328) and Korean Brain Research Institute (KBRI) basic research program through KBRI funded by the Ministry of Science and ICT (grant number: 25-BR-08-01).

## Author contributions

D.G.K. and I.S.C conceived and supervised this project. D.H.K and J.H.K developed the model and harvested the tissue needed to complete this project. S.A.C., M.T.J., K.S.K., I.Y.P. and M.Y.N. performed and analyzed biological *in vivo* experiments. S.A.C., D.H.K., M.T.J., D.G.K. and I.S.C. wrote the manuscript.

