## Supplemental Information for "Influenza A Virus infection is associated with TDP-43 pathology and neuronal damage in the brain"

### Supple Figure 1

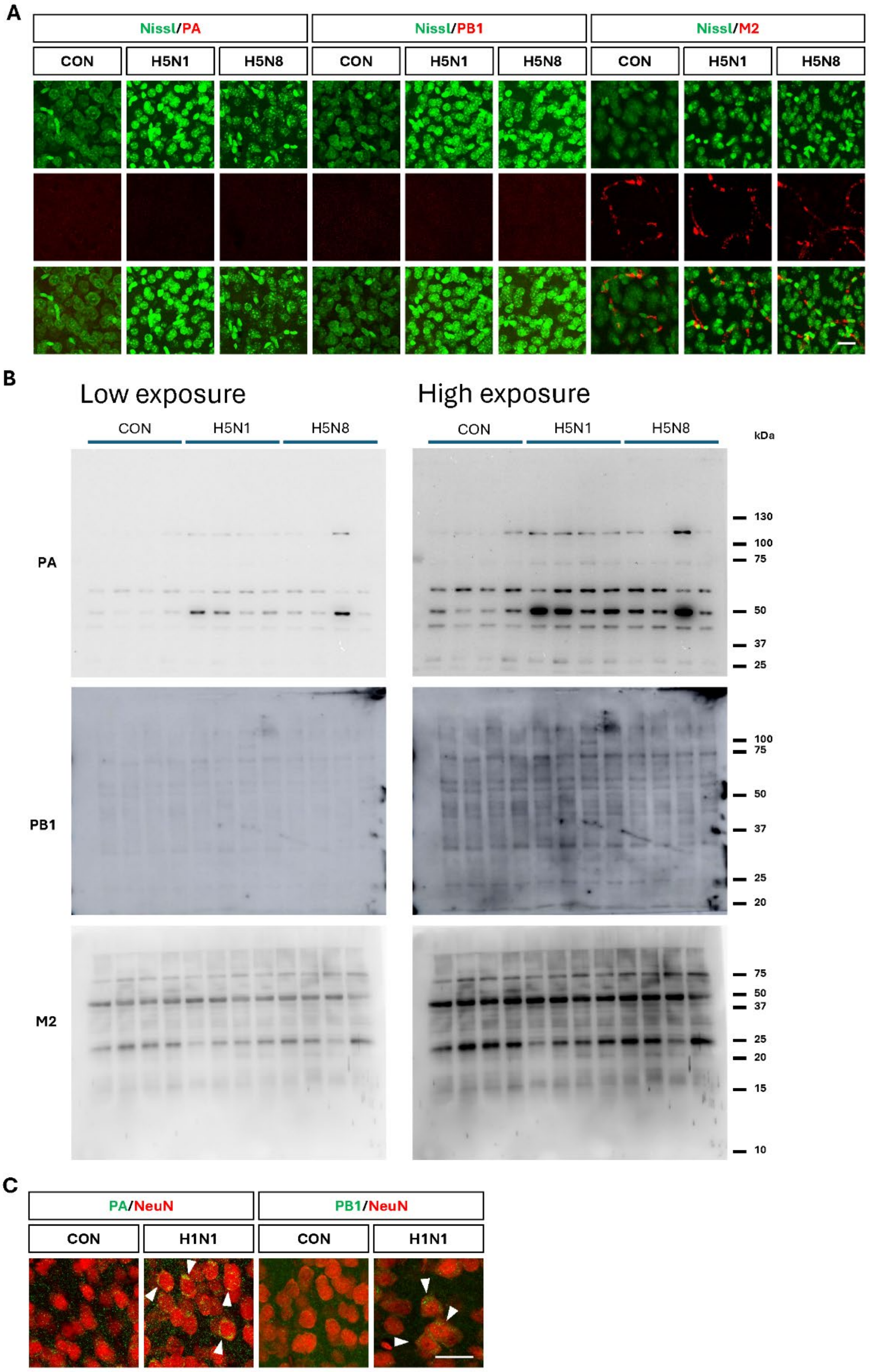

**Figure S1: Viral protein detection in the brains of IAV-infected mouse. A.** Immunostaining for influenza viral proteins, such as PA (left), PB1 (middle) and M2 (right) on the cortex of the HPAI-infected mouse brain. Nissl staining was utilized to visualize neurons. Scale bar, 20 μm. **B.** Western blotting results for viral proteins produced by Influenza A injection, PA (upper), PB1 (middle) and M2 (low), in brain tissue for three treatment groups under low and high exposure (n = 4 each groups). **C.** Immunostaining for influenza viral proteins, PA and PB1, and NeuN of the LPAI-infected mouse brain. Scale bar, 50 μm.

### Supple Figure 2

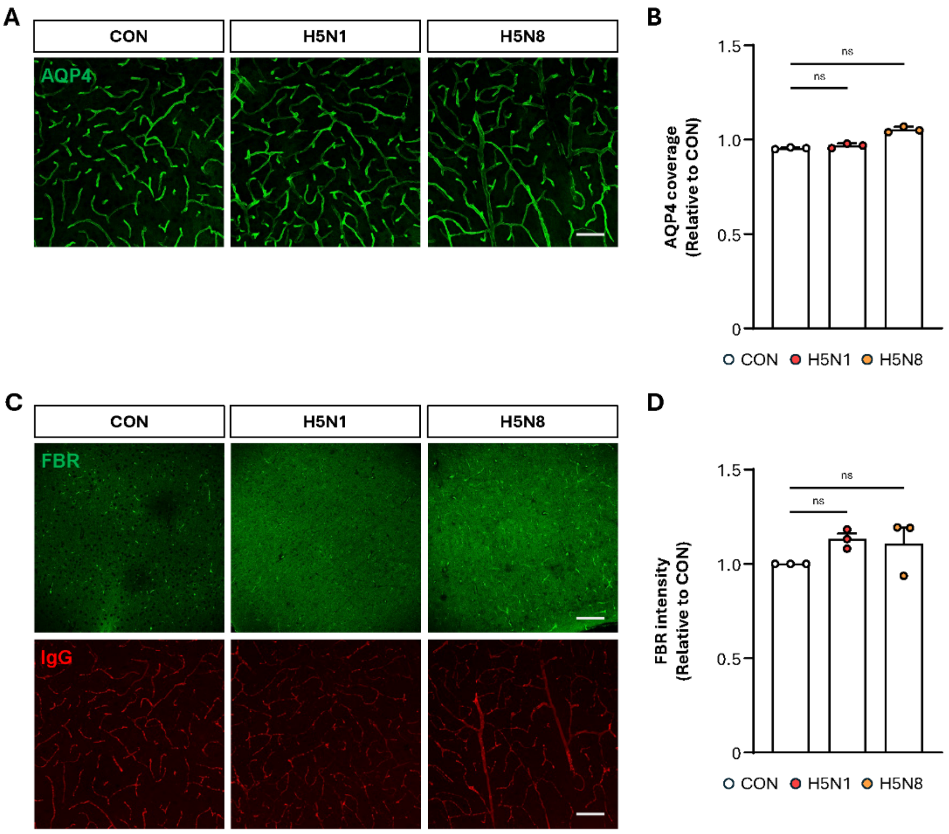

**Figure S2: Influenza A infected brain shows no alterations in BBB permeability**. **A.** Immunostaining showing neurovascular integrity through AQP4 (green) and IgG (red). Scale bar, 100 µm. **B.** Graph indicates AQP4 coverage in cortical region of brain tissue (error bars indicate means with SEM; n = 5). **C.** Immunostaining with FBR (green) indicates blood vessel integrity in the brain tissue. Scale bar, 100 µm. **D.** Graph indicates FBR intensity relative to the control in cortical region of brain tissue (error bars indicate means with SEM; n = 5).

Supple Figure 3

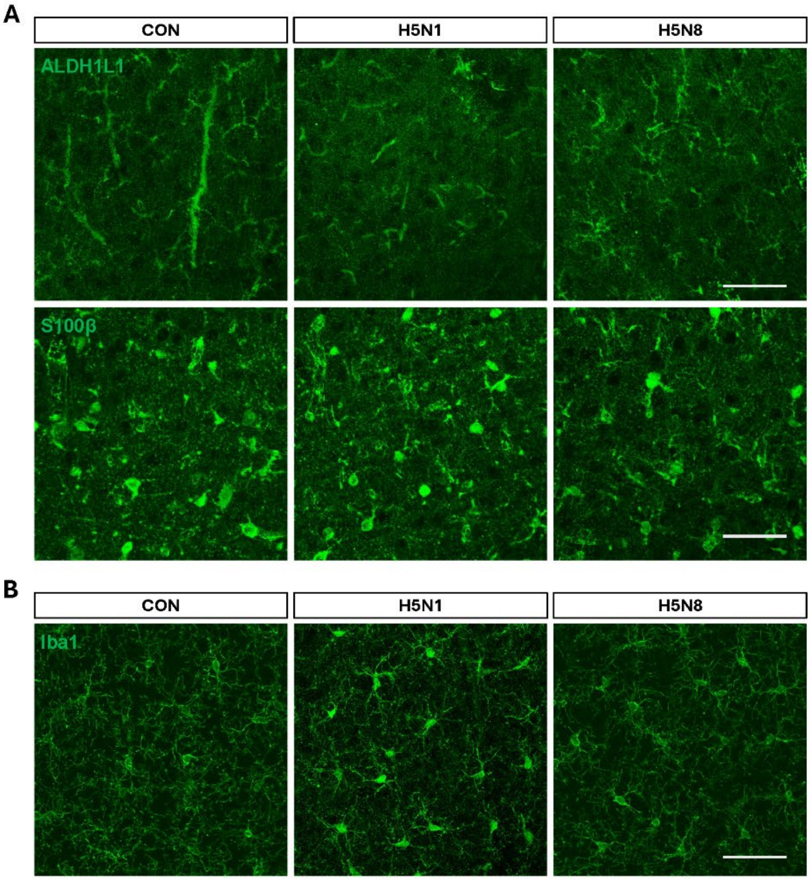

**Figure S3: Influenza A exhibits signs of astrocyte activation evasion and microglial activation under H5N1 viral infection**. **A.** Immunostaining showing astrocytic activity through ALDH1L1 (green) and S100β (green). Scale bar, 50 μm. **B.** Immunostaining showing microglial activity through Iba1. Scale bar, 50 μm.

Supple Figure 4

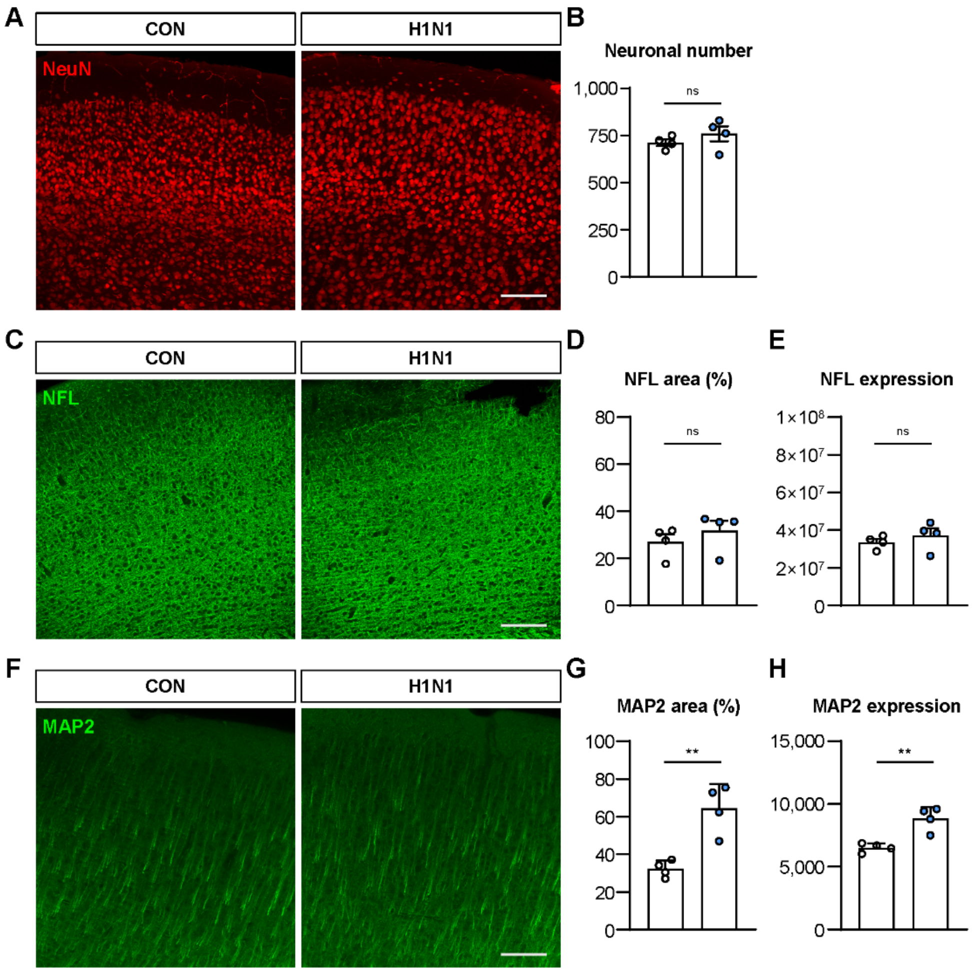

**Figure S4: Comparing neuronal effects in Influenza A positive control H1N1. A.** Immunostaining for NeuN in negative and positive control groups. Scale bar, 100 µm. **B.** Graph indicates number of NeuN-positive neurons per 500 $\mu m^2$ in the cortical region of brain tissue (error bars indicate means with SEM; n = 4). **C.** Immunostaining for NFL. Scale bar, 100 µm. **D.** Graph indicates NFL area (%) in the cortical region of brain tissue (error bars indicate means with SEM; n = 4). **E.** Graph indicates integrated density of NFL in the cortical region of brain tissue (error bars indicate means with SEM; n = 4). **F.** Immunostaining for MAP2. Scale bar, 100 µm. **G.** Graph indicates MAP2 area (%) in the cortical region of brain tissue (** indicate $p < 0.01$, determined by one-way analysis of variance (ANOVA) and Dunnet's multiple comparison's test, error bars indicate means with SEM; n = 4). **I.** Graph indicates integrated density of MAP2 in the cortical region of brain tissue (** indicate $p < 0.01$, determined by one-way analysis of variance (ANOVA) and Dunnet's multiple comparison's test, error bars indicate means with SEM; n = 4).

Supple Figure 5

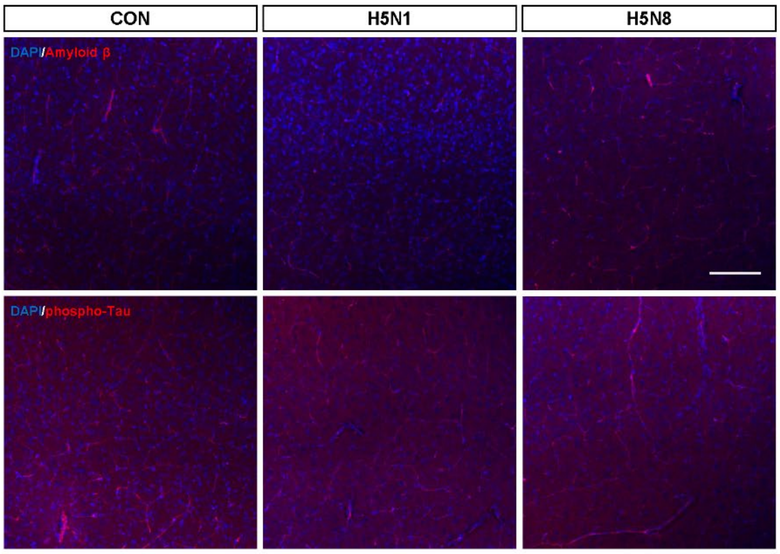

**Figure S5: Expression of pathogenic proteins involved in AD in IAV-infected mouse brain**. Immunostaining for Amyloid-β (upper) and phosphor-Tau (lower) showing that there is no changes of those proteins expression in the brains of IAV-infected mouse. Scale bar, 100 μm.

### Supple Figure 6

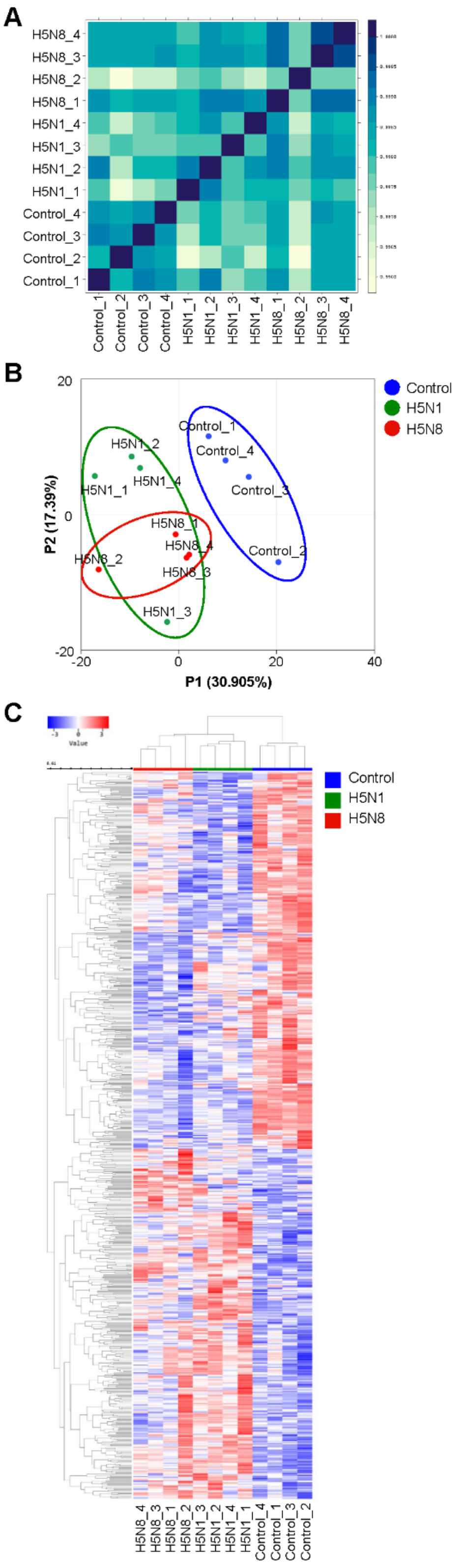

**Figure S6: Quality control data of RNA sequencing analysis. A.** The heatmap displays Pearson's correlation coefficients for all samples. **B.** The principal component analysis (PCA) plot illustrates the variability among samples, highlighting the separation between viral infection groups and the control group. **C.** The heatmap shows differentially expressed genes (DEGs) between the viral infection groups and the control groups.

**Table S1.** List of primary antibodies

| Antibody | Species | Concentration ratio | Supplied by |
|---|---|---|---|
| **ALD1H1** | Anti-rabbit | 1:500 | Abcam |
| **Influenza A M2** | Anti-mouse | 1:500 | Abcam |
| **TDP-43** | Anti-rabbit | 1:500 | Abcam |
| **Influenza A PA** | Anti-rabbit | 1:500 | ThermoFisher |
| **Influenza A PB1** | Anti-rabbit | 1:500 | ThermoFisher |
| **GFAP** | Anti-mouse | 1:500 | Cell Signaling Technology |
| **NeuN** | Anti-mouse | 1:500 | Cell Signaling Technology |
| **AQP4** | Anti-rabbit | 1:500 | Cell Signaling Technology |
| **NFL** | Anti-rabbit | 1:500 | Cell Signaling Technology |
| **Fibrinogen** | Anti-rabbit | 1:500 | Dako, Santa Clara, USA |
| **IgG** | Anti-rat | 1:500 | BioLegend |
| **CD31** | Anti-rat | 1:500 | BioLegend |
| **S100β** | Anti-mouse | 1:500 | Merck |
| **Iba1** | Anti-rabbit | 1:500 | Wako Pure Chemical Industries, Japan |
| **Amyloid β** | Anti-rabbit | 1:500 | Invitrogen |
| **Phospho-Tau** | Anti-Mouse | 1:500 | ThermoFisher |
| **TDP-43** | Anti-rabbit | 1:500 | ProteinTech, Rosemont, USA |
| **Phospho-TDP-43** | Anti-rabbit | 1:500 | ProteinTech, Rosemont, USA |
| **MAP2** | Anti-rabbit | 1:500 | Invitrogen |
| **Nissl NeuroTracer** | Alexa 488 | 1:500 | Invitrogen |
| **Fluoromyelin** | Alexa 568 | 1:500 | Invitrogen |
| **TUNEL TdT stain** | Alexa 488 | N/A | Invitrogen |
